# Poor seed quality, reduced germination, and decreased seedling vigor in soybean is linked to exposure of the maternal lines to drought stress

**DOI:** 10.1101/590059

**Authors:** Chathurika Wijewardana, K. Raja Reddy, L. Jason Krutz, Wei Gao, Nacer Bellaloui

**Author notes:** Corresponding author (KRR).

## Abstract

Effects of environmental stressors on the parent may be transmitted to the F1 generation of plants that support global food, oil, and energy production for humans and animals. This study was conducted to determine if the effects of drought stress on parental soybean plants are transmitted to the F1 generation. The germination and seedling vigor of F1 soybean whose maternal parents, Asgrow AG5332 and Progeny P5333RY, were exposed to soil moisture stress, that is, 100, 80, 60, 40, and 20% replacement of evapotranspiration (ET) during reproductive growth, were evaluated under controlled conditions. Pooled over cultivars, effects of soil moisture stress on the parents caused a reduction in the seed germination rate, maximum seed germination, and overall seedling performance in the F1 generation. The effect of soil moisture stress on the parent induced an irreversible change in the seed quality in the F1 generation and the effects on seed quality in the F1 generation were exasperated when exposed to increasing levels of drought stress. Results indicate that seed weight and storage reserve are key factors influencing germination traits and seedling growth. Our data confirm that the effects of drought stress on soybean are transferable, causing reduced germination, seedling vigor, and seed quality in the F1 generation.

## Introduction

Soybean [*Glycine max* (L.) Merr.] is a globally important annual crop, and one of the major export commodities providing oil and protein for both human and animal food. It is the second most planted field crop in the U.S. next to corn [1] and accounts for about 90% of U.S. oilseed production. Hence, soybean production provides a substantial input to the economic structure of the United States.

Soil moisture stress causes extensive losses to soybean production annually, and losses due to drought stress are projected to increase due to climate change. Climate change models indicate that historical precipitation patterns will change, and drought stress will become more severe in soybean growing regions in the United States [2]. As soil moisture stress episodes are remarkably aggravated, in recent years, greater importance has been directed to research on the unfavorable effects of soil moisture stress on soybean crop performance and yield.

It is well known that variations in environmental conditions such as photoperiod, water, nutrient status, and solar radiation can have an effect on plant growth and development [3]; however, we now understand that some effects of environmental stressors are transmittable and have fitness and phenotype costs on the F1 generation [4,5,6]. For instance, Nosalewicz et al. [7] reported that exposing barley (*Hordeum vulgare* (L.)) to drought stress during reproductive stages decreased the shoot: root ratio and the number of thick roots in the F1 generation. Moreover, exposing *Astragalus nitidiflorus* to drought stress increased seed dormancy in the F1 generation [8].

If effects of environmental stressors are transmittable to the F1 generation of plants that we depend on for food and fiber, then when, where, and how commercial seed is produced may be more critical than originally thought. We know that the quantity and quality of seed are reduced when produced in environments that have erratic precipitation patterns, high evapotranspiration demands, and high atmospheric temperatures, and that seed quality effects germination and seedling vigor [5,6,9,10]. The majority of the seed sold is produced in the same region where it is planted commercially [5].

There is growing evidence that the effects of environmental stress are transmittable to the F1 generation [4,11]. Environmental conditions influence seed size, the concentration of stress hormones in the seed, and germination rates [12,13]. Elevated concentrations of stress hormones in seed may affect the physiology and phenotypic expression through activation of the abscisic acid responsive element controlled genes [12]. Additionally, epigenetic mechanisms that affect gene activity without changing the underline DNA sequence are transmitted to successive generations leading to phenotypic modifications in the offspring [14,15]. Many studies on *Arabidopsis thaliana* indicate that stress-induced responses are inherited through the plant’s transgenerational stress memory where the offspring adaptation to particular stress is determined by the stress response established by the parent [16,17]. Some studies have also reported that previous exposure to certain stress would help the plant to acquire tolerance or to develop adaptive mechanisms to a same or different kind of stress during the crop cycle or over the generations [18,19]. To date, studies on transgenerational effects have been restricted to short-lived annual plants like *Arabidopsis thaliana*. The objective of this study was to determine if the effects of drought stress on soybean are transmittable to the F1 generation and alter seed germination and seedling vigor of the progeny in low soil moisture level environments.

## Materials and Methods

### Parental and progeny generations

Three independent experiments were conducted at the Environmental Plant Physiology Laboratory, Mississippi State University, MS, USA during the 2015 through 2017 growing seasons. The first experiment was conducted in the Soil-Plant-Atmosphere-Research (SPAR) chambers in 2015. Seed from an indeterminate, maturity group V (Asgrow AG5332) and a determinate, maturity group V soybean cultivar (Progeny P5333RY) were sown into 15.2-cm by 30-cm high polyvinylchloride (PVC) pots that contained a 3:1 mixture of a sand: loam (87% sand, 2% clay, and 11% silt). Before imposing drought stress at R1, experimental units were maintained at ambient conditions outside the SPAR units. Forty-one days after seeding (DAS) until physiological maturity, 126 DAS, the cultivars were placed in separate SPAR units and exposed to 5 different levels of drought stress, 100, 80, 60, 40, and 20% evapotranspiration (ET) replacement (Table 1) [20]. Parental lines self-fertilized under uniform SPAR conditions and generated 10 distinct F1 genetic lines, each representing a distinct cultivar by drought stress treatment. At harvest, pods were collected, air-dried at room temperature, and seeds separated manually for inclusion in a subsequent germination and vigor experiments.

**Table 1.** Analysis of variance across soybean cultivars, soil moisture stress treatments, and their interactions.

| Biomass parameter | Source |  |  |
| --- | --- | --- | --- |
|  | Treatment (Trt) | Cultivar (Cul) | Trt × Cul |
| Pods, no. plant<sup>-1</sup> | †*** | *** | NS |
| Seed, no. plant<sup>-1</sup> | *** | NS | NS |
| Seed weight, mg seed<sup>-1</sup> | *** | ** | NS |
| Yield, g plant<sup>-1</sup> | *** | ** | * |

†Results of the analysis of variance (ANOVA) indicated as \*, \*\*, \*\*\*, and NS representing significance at the $P \leq 0.05$ , $P \leq 0.01$ , $P \leq 0.001$ , and non-significant ( $P \geq 0.05$ ), respectively.

### Measurement of seed quality and chemical composition

Seeds from the first experiment were evaluated for protein, oil, fatty acids, sugars, and mineral content as previously described [21]. Briefly, seed protein and oil were determined using near infrared (NIR) spectroscopy using a diode array feed analyzer AD 7200 (Perten, Springfield, IL) at USDA research facility in Stoneville, MS, USA [22]. Perten’s Thermo Galactic Grams PLS IQ software was used to produce calibration equations, and the curves were established according to AOAC methods [23,24,25]. Analyses of fatty acids were performed on an oil basis [22,26]. Seed sugar content for glucose, sucrose, raffinose, and stachyose was determined using near-infrared reflectance (AD 7200, Perten, Springfield, IL) and analyzed based on a seed dry matter basis [22,26,27]. The concentrations of seed mineral nutrients including nitrogen (N), phosphorus (P), potassium (K), Calcium (Ca), Magnesium (Mg), Zinc (Zn), Copper (Cu), Boron (B), Iron (Fe), and Manganese (Mn) were determined at the Nutrient Testing Laboratory, Mississippi State University, MS, USA using standard procedures [28].

### Seed germination time course data for parents and offspring

To examine transgenerational effects of soil moisture stress on the F1 generation, seed germination characteristics, seedling fitness, and tolerance to osmotic stress were evaluated *in-vitro* using polyethylene glycol (PEG, molecular weight 8000 Sigma-Aldrich Company, St. Louis, MO, USA) to mimic drought stress. Using a previously described procedure, four replications of 100 seed of Asgrow AG5332, Progeny P5333RY, and the 10 F1 lines developed during experiment 1 were exposed to six levels of osmotic stress including 0.0, −0.1, −0.3, −0.5, −0.7, −0.9 [29]. The incubator was set at 25°C, which was the reported optimum temperature for the germination of soybean seed [30]. The plastic trays were vertically stacked inside the incubator and rearranged every 4 h to minimize the potential of small temperature fluctuations. Seeds were considered germinated when the radicle length was greater than half the seed length. Counts were discontinued if no seed in a tray germinated for five consecutive days.

### Curve fitting procedure and data analysis

Germination time-course was fitted with a 3-parameter sigmoidal function (Eq. 1) using SigmaPlot 13 (Systat Software Inc., San Jose, CA, USA):

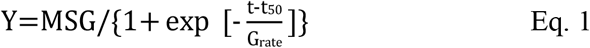

where Y is the cumulative seed germination percentage, MSG is the maximum seed germination percentage, t_50_ is the time to 50% maximum seed germination, and G_rate_ is the slope of the curve [31].

Maximum seed germination and germination rate responses to osmotic potential were analyzed using quadratic (Eq. 2) and linear (Eq. 3) regression functions. Based on the highest coefficient of determination (r^2^), MSG was modeled using a quadratic function whereas SGR was modeled by a linear function. [31]. These model functions provided regression constants to estimate maximum osmotic potential when seed germination was zero (MSGOP_max_) (Eq. 4) and maximum osmotic potential when seed germination rate was zero (SGROP_max_) (Eq. 5), correspondingly, for both the cultivars [32, 33].

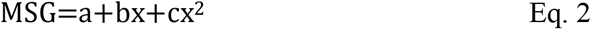

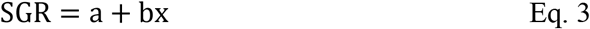

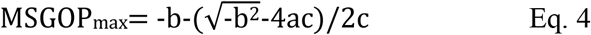

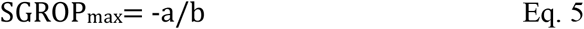

where *x* is the treatment osmotic potential and *a* and *b* are cultivar-specific equation constants generated using regression functions in SigmaPlot 13.

The effect of duration, osmotic potential, and cultivar on cumulative percent germination was tested using three-way ANOVA, followed by a Tukey multiple comparison tests. The same statistics were applied to analyze the effect of maternal soil moisture stress effects on offspring germination-based parameters and seedling growth. Also, the replicated values of SGROP_max_ and MSGOP_max_, estimated from linear and quadratic functions, were analyzed using the PROC GLM procedure of SAS. Means were compared using Fisher’s protected least significant difference at *P* < 0.05 probability. The Environmental Productivity Index (EPI) was utilized to quantify the soil moisture stress effects on estimated seed germination-based parameters such as MSG, SGR, SGROP_max_, and MSGOP_max_ [34,35,36]. To obtain soil moisture stress indices for the above parameters, the values were normalized by dividing estimated values at control treatment (100% ET) by all estimated values. The resultant relative indices ranged from 0 to 1, where, 1 indicates zero stress or the optimum soil moisture and 0 indicates total stress or severe water deficit. The developed EPI values were tested to understand the difference between the cultivars and the regression analyses were performed on the relationship between derived values and soil moisture. Critical limits for seed germination-based parameters as a function of soil moisture content were calculated as 90% of the control of soil moisture treatment.

### Maternal effects on seedling vigor

To examine the transgenerational effects of soil moisture stress on seed emergence and seedling vigor, the F1 generation from experiment 2 was exposed to three levels of drought stress, 100%, 66%, and 33% of field capacity. The experiment was conducted using pre-fabricated mini-hoop structures (rain-out shelters). Each structure consisted of a PVC framework with 4 MIL polythene wrapping having the dimensions of 2-m width × 1.5-m height × 5-m length. Seeds were sown in PVC pots (15.2-cm diameter by 30.5-m high) that contained a 3:1 mixture of sand: loam (87% sand, 2% clay, and 11% silt). Pots were arranged in a completely randomized design with 4 replications per cultivar organized in 2 rows with twenty-four pots per row. Four seeds were sown in each pot but were thinned to one per pot 1 week after emergence. From emergence until 6 DAS, plants were maintained at 100% of FC. Drought stress treatments were imposed 7 DAS and continued until harvest, 29 DAS. Throughout the experimental period, plants were fertigated with full-strength Hoagland’s nutrient solution delivered through an automated and computer controlled drip irrigation system. Soil moisture contents were monitored using Decagon soil moisture sensors (5TM Soil Moisture and Temperature Sensor, Decagon Devices, Inc., Pullman, WA) inserted at a depth of 15 cm in five random pots of each treatment.

### Measurements

#### Physiological and gas-exchange measurements

AT 26 DAS, leaf chlorophyll (Chl) content, epidermal flavonoids (Flav), epidermal anthocyanin (Anth), and nitrogen balance index (NBI) were measured on the uppermost recently fully expanded leaf with a Dualex^®^ Scientific Polyphenols and Chlorophyll Meter (FORCE-A, Orsay, France). Gas exchange and chlorophyll fluorescence parameters were measured using the LI-6400 photosynthesis system (LiCOR Inc., Lincoln, NE). Measurements were made on the uppermost recently fully expanded leaf from three plants in each cultivar from each treatment between 1000 and 1200 h. While measuring photosynthesis (Pn), the instrument was set at 1500 μmol photon m^−2^ s^−1^ photosynthetically active radiation, daytime temperature 29°C, 410 μmol mol^−1^ CO_2_, and 50% relative humidity. The flow rate through the chamber was adjusted to 350 mol s^−1^. Pn and the fluorescence (Fv/Fm) were recorded as the total coefficient of variation (%CV) reached a value of less than 0.5. The instrument itself calculates stomatal conductance (gs), transpiration (E), and electron transport rate (ETR) by considering incoming and outgoing flow rates and leaf area. Intrinsic water use efficiency (WUE) and the ratio of internal (C_i_) to external (C_a_) CO_2_ concentration were estimated as the ratio of Pn/Trans and C_i_/C_a_.

### Growth and biomass components

At harvest plant height and the number of nodes were recorded. Leaf area was measured using the LI-3100 leaf area meter (LI-COR, Inc., Lincoln, NE). Stems, leaves, and roots were separated from each plant, and total dry weight per plant was calculated by adding the dry weight of different plant components after oven drying at 80°C for 5 days.

### Root morphology

After separating the stem from the root system of each plant, roots were washed with water on a sieve [37,38,39]. Washed roots were scanned with a WinRhizo Pro optical scanner (Regent Instruments, Inc., QC, Canada) by floating the individual root system in 5 mm of water in a Plexiglas tray [39,40]. Gray-scale root images were acquired by setting the parameters to high accuracy (resolution 800 × 800 dpi), and the images were analyzed using WinRhizo Pro software. From the scanned images, WinRhizo Pro software calculated seven root parameters including, root length, surface area, diameter, volume, number of tips, forks, and crossings.

### Data analysis

Analysis of variance (ANOVA) was performed to determine the differences among the cultivars and treatments for the seed quality traits, mineral composition, growth, physiological, and root developmental parameters using SAS 9.2 (SAS Institute 2011, Cary, NC). Multiple comparison analyses were performed to identify cultivar specific responses to treatment effects at the *P* = 0.05 level of significance using *t*-test. Graphical analysis was carried out using Sigma Plot 13.0 (Systat Software Inc., San Jose, CA).

## Results and Discussion

### Seed number and individual seed weight

Soil moisture stress during reproductive stages had an adverse effect on soybean pod and seed number, individual seed weight, and seed yield (Table 2, Fig 1). Pooled over cultivar, seed number was positively correlated with soil moisture level. As soil moisture stress intensified, not only did the number of seed decrease, but the production of small, shriveled, and wrinkled seeds in the seed lot increased (Fig 2). At severe water deficit (20% ET), most of the seeds were long-oval shaped and shrunken, whereas, under well-irrigated condition (100% ET), seeds were large, full, and rounded in shape. Others noted that soil moisture stress caused a reduction in seed number and individual seed weight due to the reduction in biomass accumulation and partitioning towards seeds when plants were under drought stress [41,42]. When soil moisture stress begins in the early reproductive stages, it causes flower abortion, while drought stress during seed fill reduces carbon assimilation and causes smaller seeds, lower seed weights, and reduced seed quality [41].

**Table 2.** Analysis of variance for different soybean seed germination based parameters.

| Source | CSG | MSG | t<sub>50</sub> | SGR |
| --- | --- | --- | --- | --- |
| Osmotic potential (Trt) | ¶*** | *** | *** | *** |
| Cultivar (Cul) | *** | *** | *** | *** |
| Time | *** | - | - | - |
| Trt × Cul | *** | *** | *** | *** |
| Cul × Time | *** | - | - | - |
| Trt × time | *** | - | - | - |
| Trt × Cul × Time | NS | - | - | - |

¶\*\*\* and NS represent significance level at $P \leq 0.001$ and $P > 0.05$ . The dash (‘-’) sign indicates that the ANOVA or the given term was not assessed. Osmotic stress treatments (Trt), soybean cultivars (Cul), and their interactions (Cul × Trt) with cumulative percent germination (CSG), maximum seed germination (MSG), time to 50% germination (t<sub>50</sub>), and seed germination rate (SGR).

**Fig 1.**
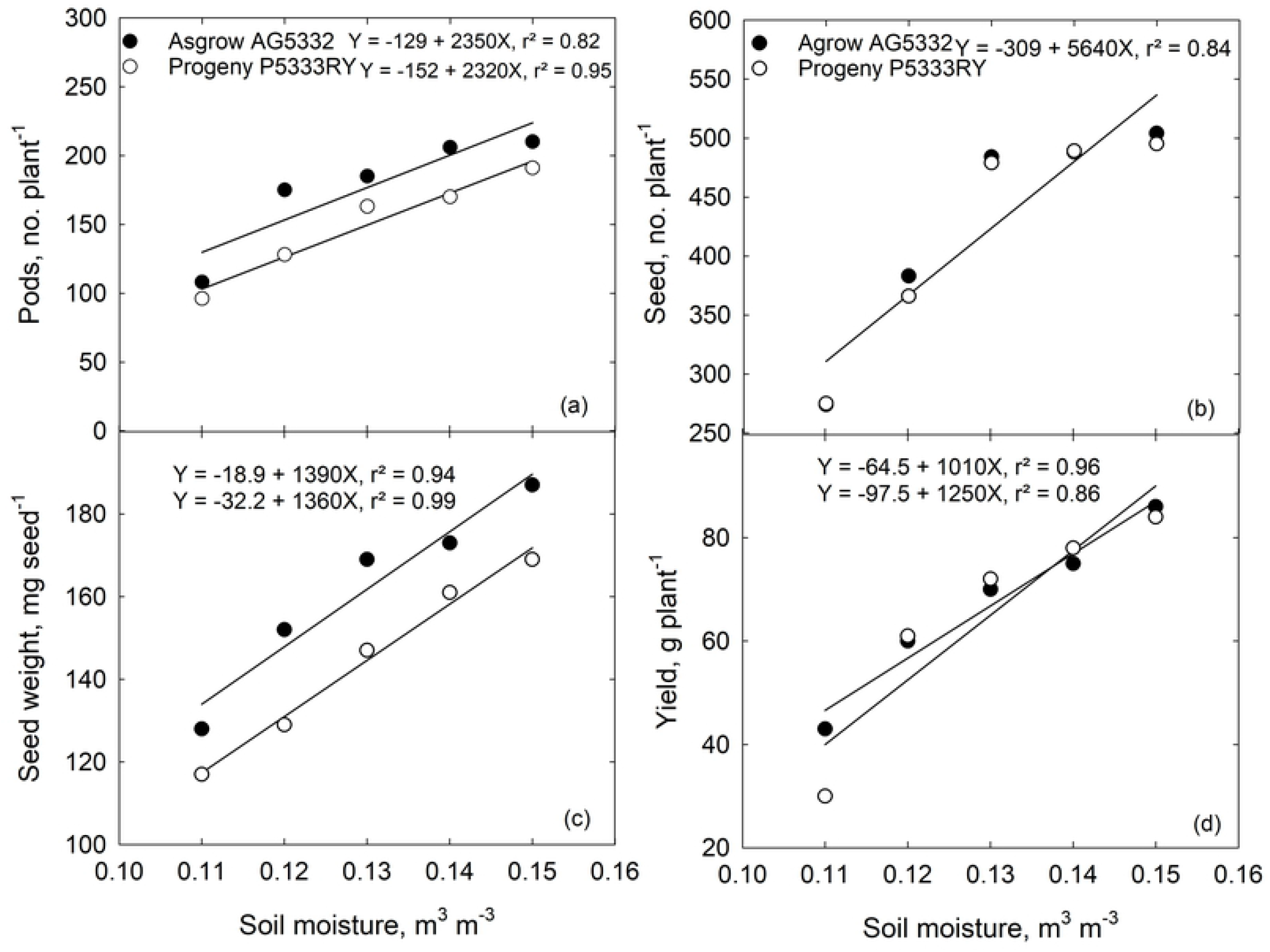
The effect of increasing soil moisture stress (100, 80, 60, 40, and 20% ET) on soybean (a) pod number, (b) seed number, (c) individual seed weight, and (d) yield of Asgrow AG5332 and Progeny P5333RY harvested 126 d after planting.

**Fig 2.**
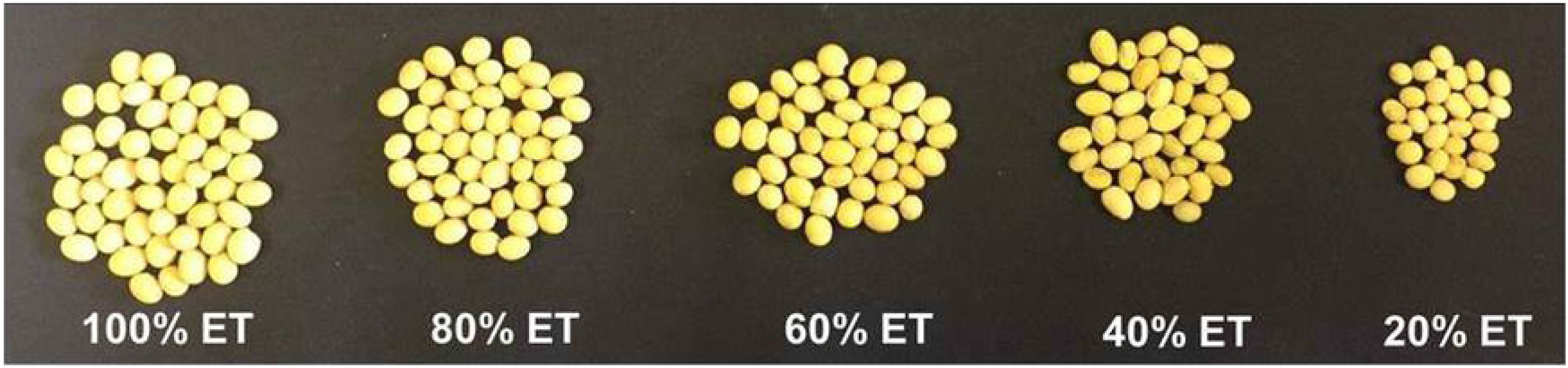
The effect of increasing soil moisture stress (100, 80, 60, 40, and 20% ET) on a soybean seed number, size, and shape of Asgrow AG5332 harvested 126 d after planting. Each seed lot under each treatment represents 1/10 of the total fraction of seeds collected from one plant. Due to the absence of significant difference for the seed number between the cultivars, seeds only from one soybean cultivar are presented.

## Seed germination traits

### Cumulative percent germination

A primary hypothesis of this research is that the effects of drought stress are transferable to the F1 generation and that the transferable effects on the F1 generation are exasperated in low osmotic potential environments. In agreement with our hypothesis, for both cultivars, seeds from well-irrigated plants (100% ET) and their parental seeds germinated more successfully and faster than seed formed under drought stressed environments (Fig 3 and 4). For a given osmotic water potential and cultivar, percent germination was inversely correlated with ET replacement level of the parental line. In general, the percent germination of Progeny P5333RY (Fig 4) was lower than that of Asgrow AG5332 (Fig 3) at all osmotic potentials except that of −0.9 MPa. No seeds from the parental lines of Progeny P5333RY exposed to 40% ET replacement or Progeny P5333RY exposed to 20% ET replacement germinated at an osmotic potential of −0.9 MPa. No Asgrow AG5332 lines germinated at an osmotic potential of −0.9 MPa. Our data indicate that the maternal environment has a strong influence on both the timing of germination and viability of soybean seeds. Seeds from stressful maternal environments, especially from 40 and 20% ET replacement, germinated later and the proportion of non-germinated seeds remained higher at the end of the experiment. Others have noted a variation in germination timing in different plant species concerning adverse maternal environments [6,11,43].

**Fig 3.**
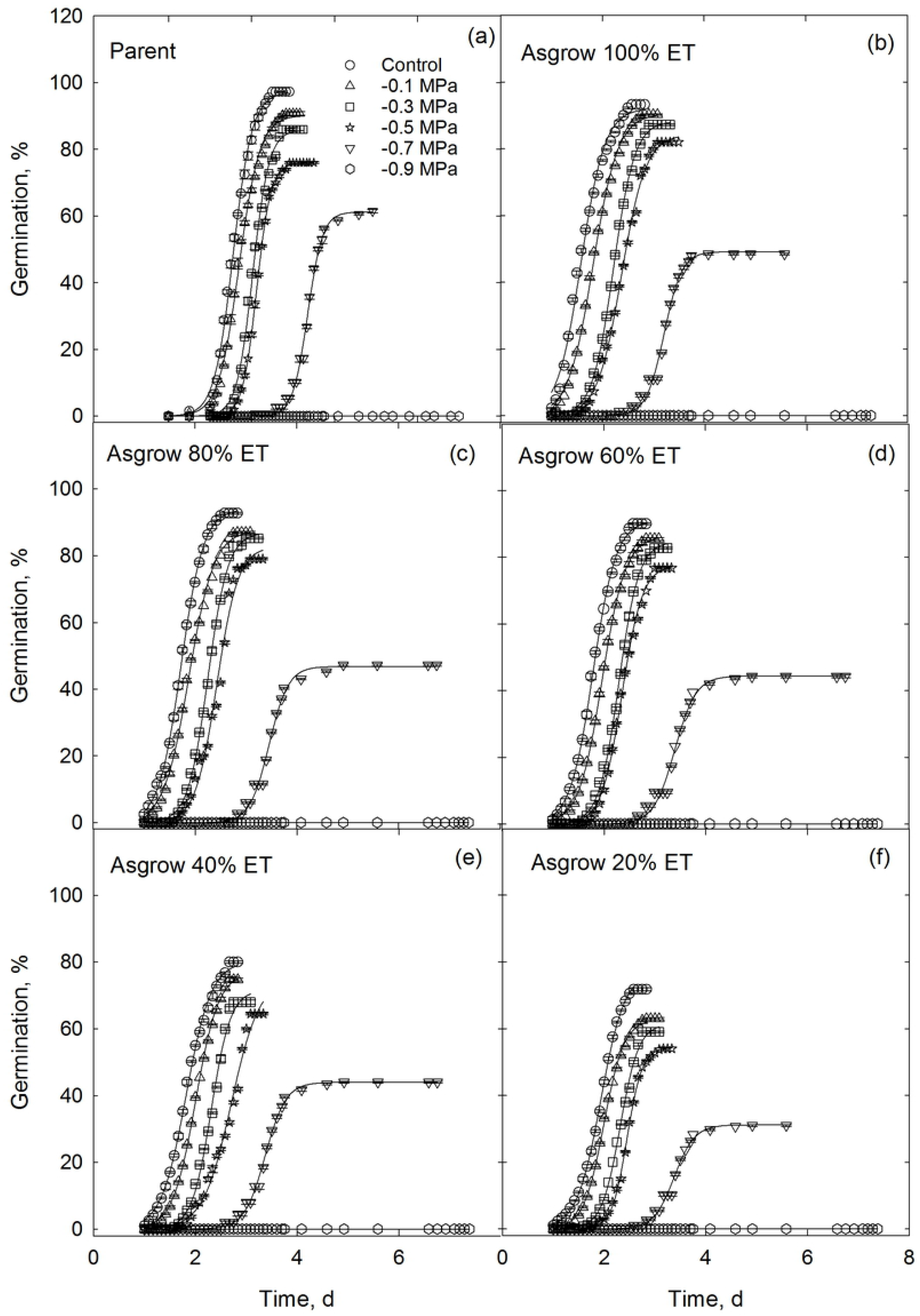
Germination of the F1 generation in osmotic potentials ranging from 0.0 to −0.9 MPa. Parent lines for the F1 seed were collected from the soybean cultivar Asgrow AG5332 after exposure to soil moisture stress including replacement of 100, 80, 60, 40, and 20% of the evapotranspiration demand. Symbols represent the mean of four replications, while bars represent the standard error. Lines are the best fit of the 3-parameter sigmoidal function.

**Fig 4.**
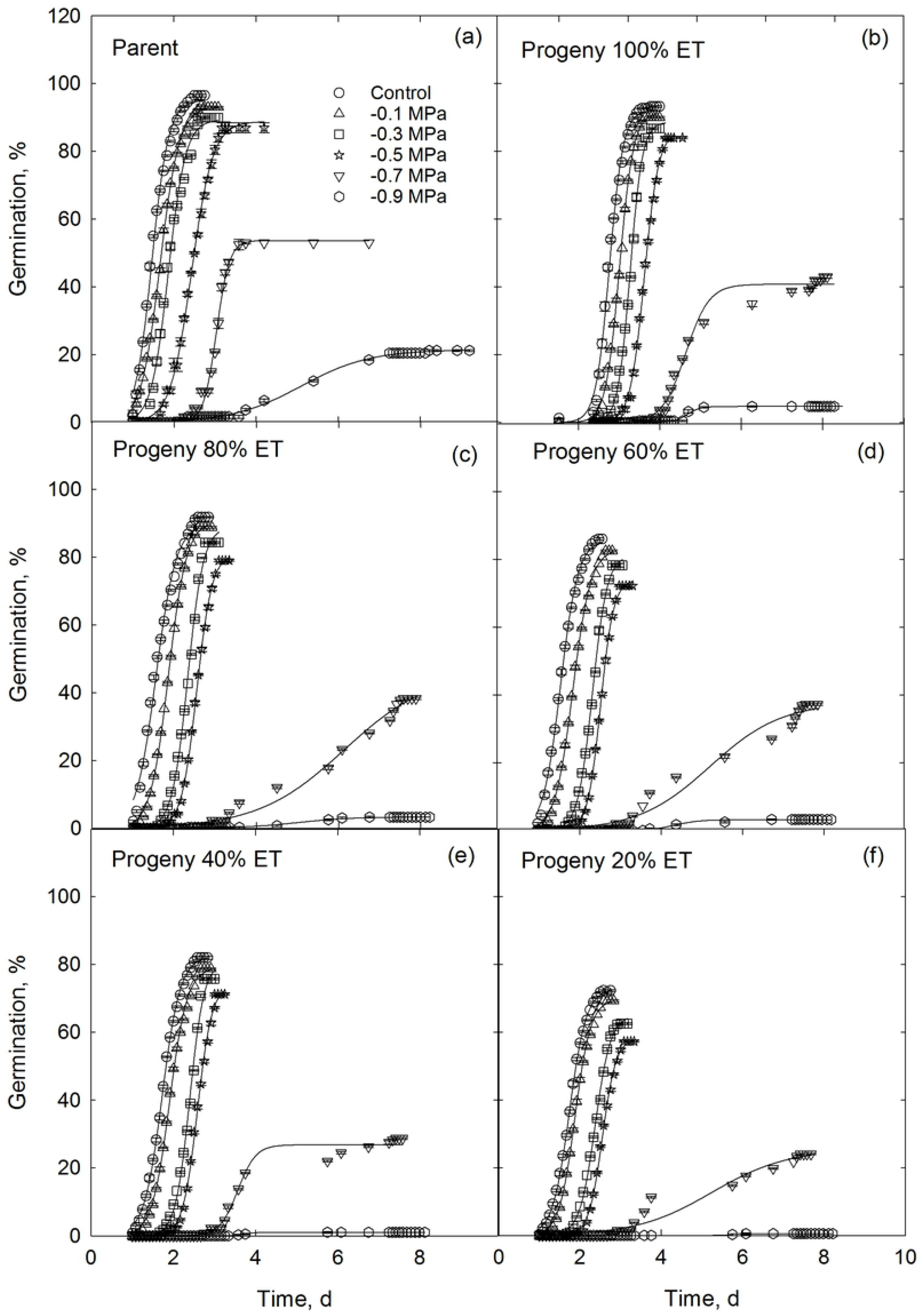
Germination of the F1 generation in osmotic potentials ranging from 0.0 to −0.9 MPa. Parent lines for the F1 seed were collected from the soybean cultivar Progeny P5333RY after exposure to soil moisture stress including replacement of 100, 80, 60, 40, and 20% of the evapotranspiration demand. Symbols represent the mean of four replications, while bars represent the standard error. Lines are the best fit of the 3-parameter sigmoidal function.

### Maximum seed germination and rate

Similar to cumulative percent germination, drought stress during the reproductive growth stages of the maternal line caused a decrease in the maximum seed germination and germination rate in the F1 generation (Table 3). Moreover, maximum germination for most osmotic potentials was inversely correlated with parents stress level at the time of seed formation (Fig 5). Seed germination rate was correlated with osmotic stress, and the relationship was best described by a linear regression model (mean r^2^ = 0.95) (Fig 6). The effect of osmotic potential on maximum germination and the rate of germination could be due to reduced imbibition of water and subsequent effects on embryo growth and development.

**Table 3.** Analysis of variance for different soybean seed germination based parameters based on the parental environment.

| Source | CSG | MSG | t<sub>50</sub> | SGR |
| --- | --- | --- | --- | --- |
| Osmotic potential (Trt) | ¶*** | *** | *** | *** |
| PE | *** | *** | *** | *** |
| Offspring (Cul) | *** | *** | ** | ** |
| Trt × PE | *** | *** | *** | *** |
| Cul × PE | *** | ** | ** | ** |
| Trt × Cul | *** | *** | ** | ** |
| Trt × Cul × PE | * | * | * | * |

¶\*\*\*, \*\*, \* and NS represent significance level at $P \leq 0.001$ , $P \leq 0.05$ , $P \leq 0.01$ , and $P > 0.05$ . Osmotic stress treatments (Trt), parental environment (PE), soybean offspring (Cul), and their interactions (Cul × Trt × PE) with cumulative percent germination (CSG), maximum seed germination (MSG), time to 50% germination (t<sub>50</sub>), and seed germination rate (SGR).

**Fig 5.**
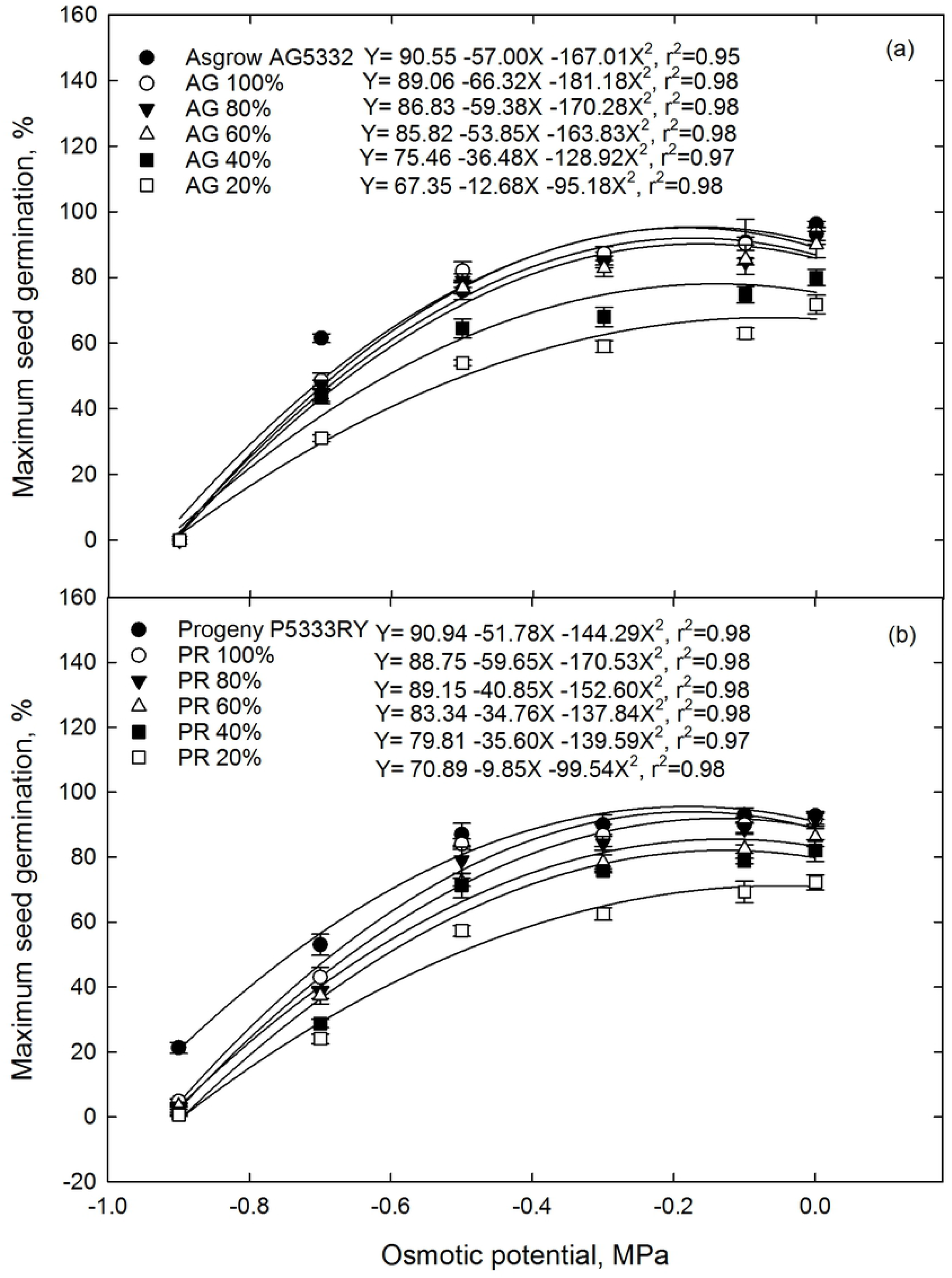
Maximum germination of soybean seed from the F1 generation as a function of osmotic potential. F1 seed was collected from the soybean cultivar (a) Asgrow AG5332 and (b) Progeny P5333RY after exposure to drought stress including replacement of 100, 80, 60, 40, and 20% of the evapotranspiration demand. Symbols represent the mean of four replications, while bars represent the standard error. Lines are the best fit of a quadratic function.

**Fig 6.**
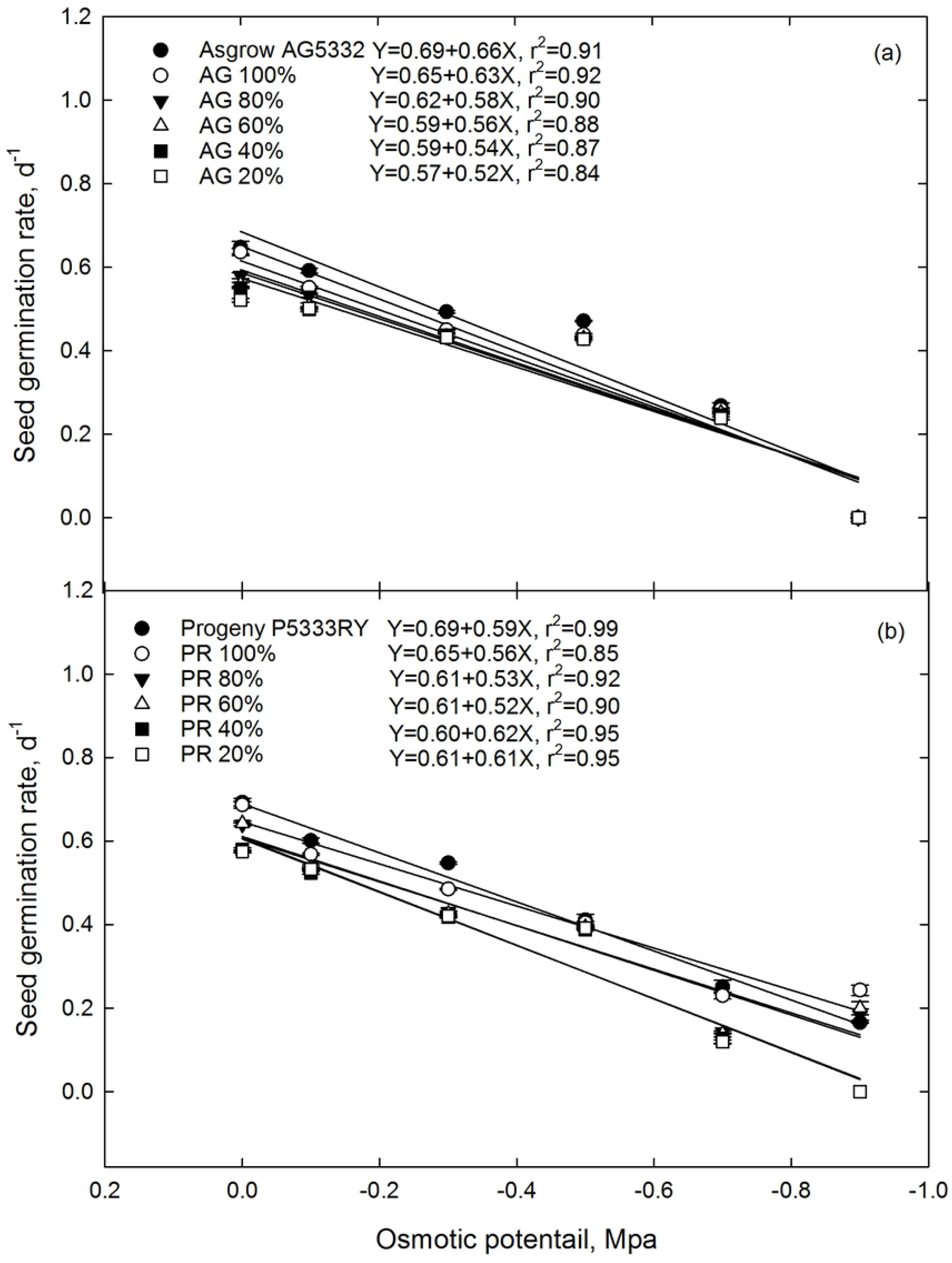
The germination rate of soybean seed from the F1 generation as a function of osmotic potential. F1 seed was collected from the soybean cultivar (a) Asgrow AG5332 and (b) Progeny P5333RY after exposure to drought stress including replacement of 100, 80, 60, 40, and 20% of the evapotranspiration demand. Symbols represent the mean of four replications, while bars represent the standard error. Lines are the best fit of a quadratic function.

The transgenerational effect of drought stress on MSG and SGR differed between cultivars and was exasperated when the F1 generation was exposed to increased osmotic potentials (Table 3). The seeds from 20% ET maternal environment showed the lowest MSG for both Asgrow AG5332 (Fig 5a) and Progeny P5333RY (Fig 5b) cultivars, where they showed 26 and 22% reduction in MSG compared to their parents. The rate of decline was greater for Progeny P5333RY relative to Asgrow AG5332 as osmotic potential increased from 0.0 to 0.9 MPa.There were transgenerational differences in seed germination parameters for cultivar and parental environments (Table 3). At an osmotic potential of 0.0 MPA, the MSG of parent line for Asgrow AG5332 (Fig 5a) was 3.3% greater than that of parent line of Progeny 5333RY (Fig 5b). Stressful maternal environments (20% ET), decreased the rate of germination in which it ranged from 0.57 d^−1^ to 0.61 d^−1^ for Asgrow AG5332 (Fig 6a) and Progeny 5333RY(Fig 6b), respectively at 0.0 MPa osmotic potential compared to their parents. Seed germination rate is a key factor which determines seed’s survival potential under desired or stressful conditions. With rapid germination, the chance of survival is much greater regarding successful stand establishment and rapid resource exploration.

### Parameter estimates

In our study, the parameter estimates, maximum osmotic potential when MSG was zero (MSGOP_max_) and maximum osmotic potential when SGR was zero (SGROP_max_) were obtained from the regression constants derived by the quadratic and linear model functions of MSG and SGR (Table 4). These values indicate cultivar’s critical germination potential at the given osmotic stress. The knowledge of seed lethal osmotic potentials where we would expect zero MSG and SGR is essential for seed conservation under limited water conditions as well as during drying and storage. According to our observations, the parameter estimates were also modified by the maternal environment (Table 4).

**Table 4.**
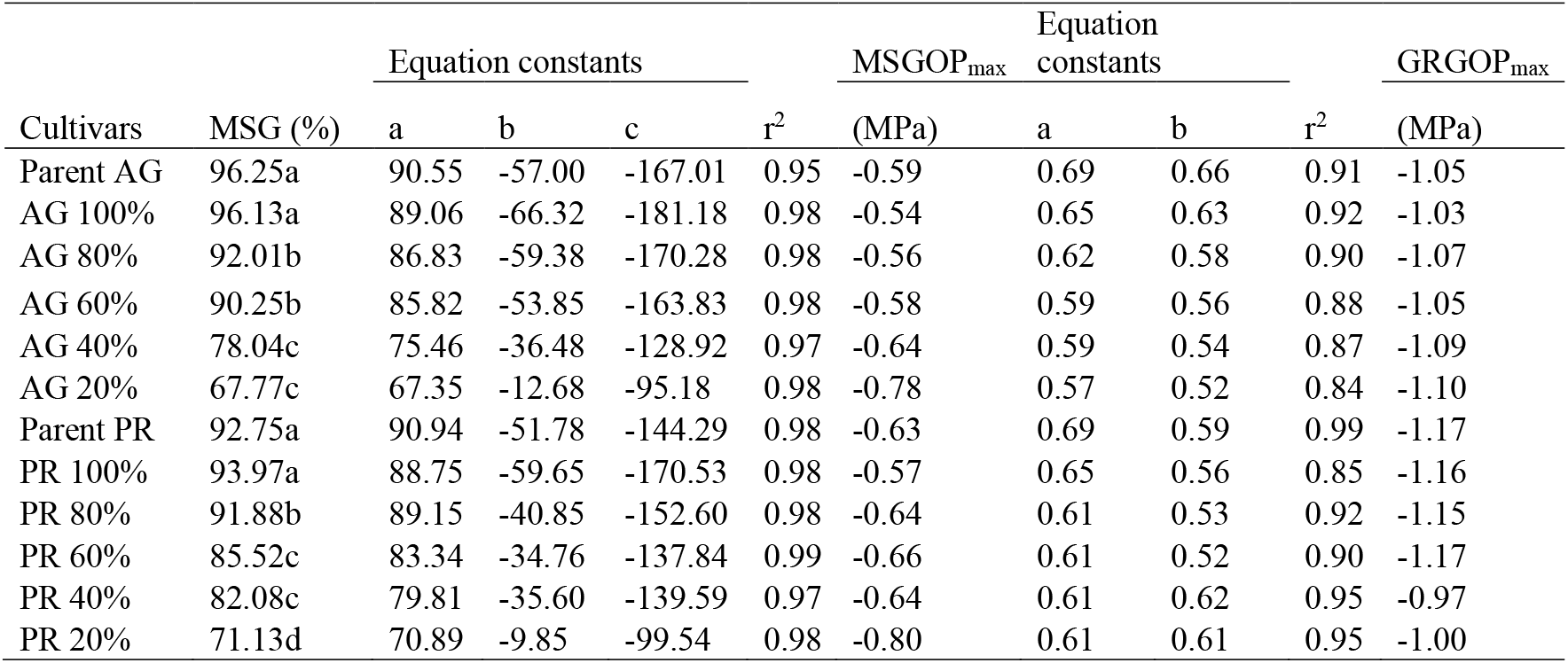
Maximum germination and estimated parameters of soybean seed from the F1 generation as a function of osmotic potential. F1 seed was collected from the soybean cultivar Asgrow AG5332 and Progeny P5333RY after exposure to drought stress including replacement of 100, 80, 60, 40, and 20% of the evapotranspiration demand. Maximum seed germination (MSG), quadratic equation constants (a, b, c), coefficient of determination (r^2^) for MSG, estimated maximum osmotic potential when seed germination was zero (MSGOP_max_), linear regression constants (a, b), coefficient of determination (r^2^) for seed germination rate, and estimated maximum osmotic potential when SGR was zero (SGROP_max_). AG and PG represent Asgrow AG5332 and Progeny P5333RY correspondingly.

| Cultivars | MSG (%) | Equation constants | | | $r^2$ | MSGOP<sub>max</sub><br>(MPa) | Equation constants | | $r^2$ | GRGOP<sub>max</sub><br>(MPa) |
| --- | --- | --- | --- | --- | --- | --- | --- | --- | --- | --- |
|  |  | a | b | c |  |  | a | b |  |  |
| Parent AG | 96.25a | 90.55 | -57.00 | -167.01 | 0.95 | -0.59 | 0.69 | 0.66 | 0.91 | -1.05 |
| AG 100% | 96.13a | 89.06 | -66.32 | -181.18 | 0.98 | -0.54 | 0.65 | 0.63 | 0.92 | -1.03 |
| AG 80% | 92.01b | 86.83 | -59.38 | -170.28 | 0.98 | -0.56 | 0.62 | 0.58 | 0.90 | -1.07 |
| AG 60% | 90.25b | 85.82 | -53.85 | -163.83 | 0.98 | -0.58 | 0.59 | 0.56 | 0.88 | -1.05 |
| AG 40% | 78.04c | 75.46 | -36.48 | -128.92 | 0.97 | -0.64 | 0.59 | 0.54 | 0.87 | -1.09 |
| AG 20% | 67.77c | 67.35 | -12.68 | -95.18 | 0.98 | -0.78 | 0.57 | 0.52 | 0.84 | -1.10 |
| Parent PR | 92.75a | 90.94 | -51.78 | -144.29 | 0.98 | -0.63 | 0.69 | 0.59 | 0.99 | -1.17 |
| PR 100% | 93.97a | 88.75 | -59.65 | -170.53 | 0.98 | -0.57 | 0.65 | 0.56 | 0.85 | -1.16 |
| PR 80% | 91.88b | 89.15 | -40.85 | -152.60 | 0.98 | -0.64 | 0.61 | 0.53 | 0.92 | -1.15 |
| PR 60% | 85.52c | 83.34 | -34.76 | -137.84 | 0.99 | -0.66 | 0.61 | 0.52 | 0.90 | -1.17 |
| PR 40% | 82.08c | 79.81 | -35.60 | -139.59 | 0.97 | -0.64 | 0.61 | 0.62 | 0.95 | -0.97 |
| PR 20% | 71.13d | 70.89 | -9.85 | -99.54 | 0.98 | -0.80 | 0.61 | 0.61 | 0.95 | -1.00 |

### Seed size, individual seed weight, and quality

Seed size and weight are key characters and known to be strongly influenced by the maternal environment. Previous studies have found that seed weight was the driving factor on the differences in germination timing due to changes in maternal environments [44]. Agreeing with our observations, individual seed weight linearly correlated with maximum seed germination (Fig 7a); with heavier and bigger seeds tending to germinate much earlier and faster. The two soybean cultivars showed significant differences (*P*<0.05) for seed weight, seed protein (Fig 7b), palmitic acid (Fig 7c), and nitrogen (Fig 7e) except sucrose (Fig 7d) and phosphorus (Fig 7f). Overall, Asgrow AG 5332 showed higher seed weight, proteins, fatty acids, sucrose, and minerals signifying the genetic variation in the sensitivity to the soybean maternal environmental variations. Similar to seed weight, seed protein, palmitic acid, sucrose, nitrogen, and phosphorus also exhibited linear correlations concerning maximum seed germination. This implies that heavier soybean seeds with the higher content of seed proteins, fatty acids, sugars, and minerals and storage reserves were able to provide energy more rapidly to germinating seed, which in turns increased the maximum seed germination and germination rate. Therefore, the parental environment during seed development apparently could have a significant impact on carbohydrate reserve, proteins, minerals, and overall seed quality.

**Fig 7.**
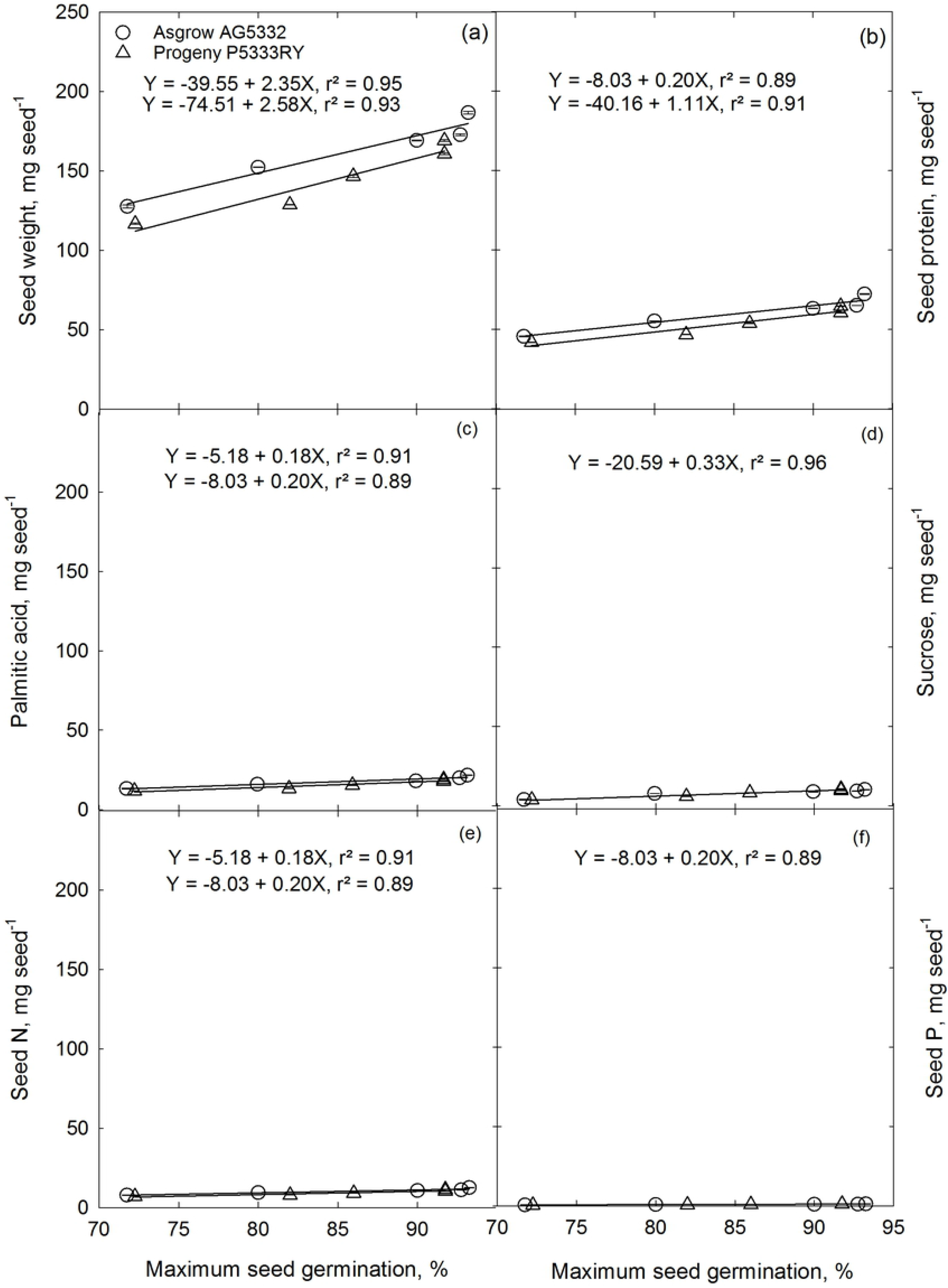
Correlation between maximum germination of soybean seed from the F1 generation of the Asgrow AG5332 and Progeny P5333RY exposed to drought stress including replacement of 100, 50, 60, 40, and 20% of the evapotranspiration demand and (a) seed weight, (b) seed protein content, (c) seed palmitic acid concentration, (d) seed sucrose concentration, (e) seed nitrogen content, and (f) seed phosphorus content. Soil moisture stress treatments were imposed on the parents at 41 days after seeding, R1 growth stage, and continued until harvest, 126 d after seeding. Symbols represent the mean of four replications. Standard error bars are not displayed when smaller than the symbol for the mean.

Although seed weight and quality appeared to affect seed germination traits for the seeds came from stressful maternal environments, some other mechanisms might have involved in the transmission of the observed maternal effects on the progeny. These include epigenetic mechanisms such as histone modifications, DNA methylation, changes in the frequency of homologous recombination, and changes in small and micro RNAs [14,16,17]. Furthermore, direct soil moisture stress effects on the accumulation of metabolites and mRNA or proteins in the seeds [4] can also play a vital role in transmitting maternal effects to the offspring. However, our experiment methodology does not permit to differentiate whether the observed changes are due to heritable or non-heritable transgenerational effects which decreased the fitness of the progeny under the water-stressed condition similar to their maternal environments.

The calculated environmental productivity indices (EPI) represented the fractional limitation due to soil moisture stress and ranged from 0 to 1 where 0 is when the soil moisture is limiting that particular seed germination trait, and 1, when it does not limit that parameter. By doing such, the effects of soil moisture on maximum seed germination, seed germination rate, and other parameter estimates such as maximum osmotic potential when MSG was zero (MSGOP_max_) and maximum osmotic potential when SGR was zero (SGROP_max_) could be quantified in a changing soil moisture environment without the other confounding factors such as temperature. In the present study, all the parameters declined linearly under moderate and severe water-stressed conditions except SGROP_max_ (Fig 8). Since the two cultivars did not show a significant difference, one linear regression was fitted for the two soybean cultivars. Based on the estimated critical limits (CL), the parameter estimates (MSGOP max CL= 1.116 m^3^ m^−3^ and SGROP_max_ CL=1.110 m^3^ m^−3^) were lower than MSG and SGR, indicating their less sensitivity towards soil moisture. The critical limit of SGR (0.139 m^3^ m^−3^) was higher than that of MSG (0.133 m^3^ m^−3^); therefore, MSG was more sensitive to soil moisture stress than SGR. This finding suggests that like 90% of optimum soil moisture condition, the critical soil moisture level would be 0.11 and 0.14 m^3^ m^−3^ for soybean maximum seed germination and germination rate.

**Fig 8.**
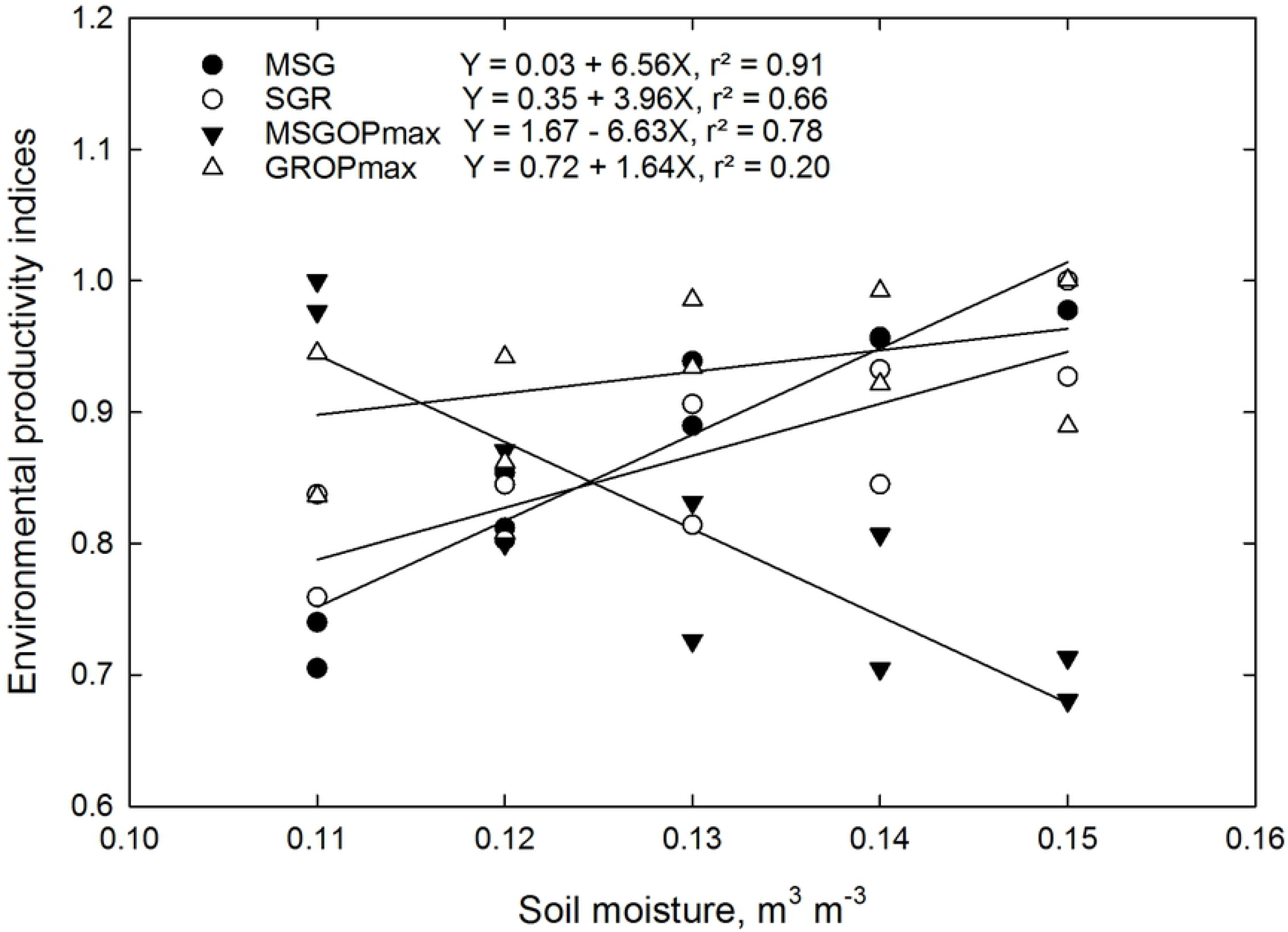
Soil moisture dependent environmental productivity indices (EPI) for maximum seed germination (MSG), seed germination rate (SGR), maximum osmotic potential when MSG was zero (MSGOP_max_), and maximum osmotic potential when SGR was zero (SGROP_max_). Potential values were estimated by dividing the measured parameters by its estimated maximum value at optimum (0.15 m^3^ m^−3^) level and expressed as a fraction between 0 and 1.

### Maternal effects on seedling establishment

A principal hypothesis of this experiment was that drought stress imposed during the reproductive stages of soybean has a transferable effect on seedling growth and development in the F1 generation. The maternal environment affected the rate of emergence of F1 seedlings (Table 5), and the effect was exasperated as the average soil moisture content (Fig 9) decreased from 0.14 to 0.07 m^3^ m^3^ (Fig. 10). Pooled over cultivar, seeds from a maternal environment receiving 100% ET replacement emerged 120% faster than seeds from a maternal environment receiving 20% ET replacement. The slower rate of emergence from soybean seeds that developed in a 20% ET replacement environment might be due to the effect of drought stress on seed mass and quality [9,42].

**Table 5.**
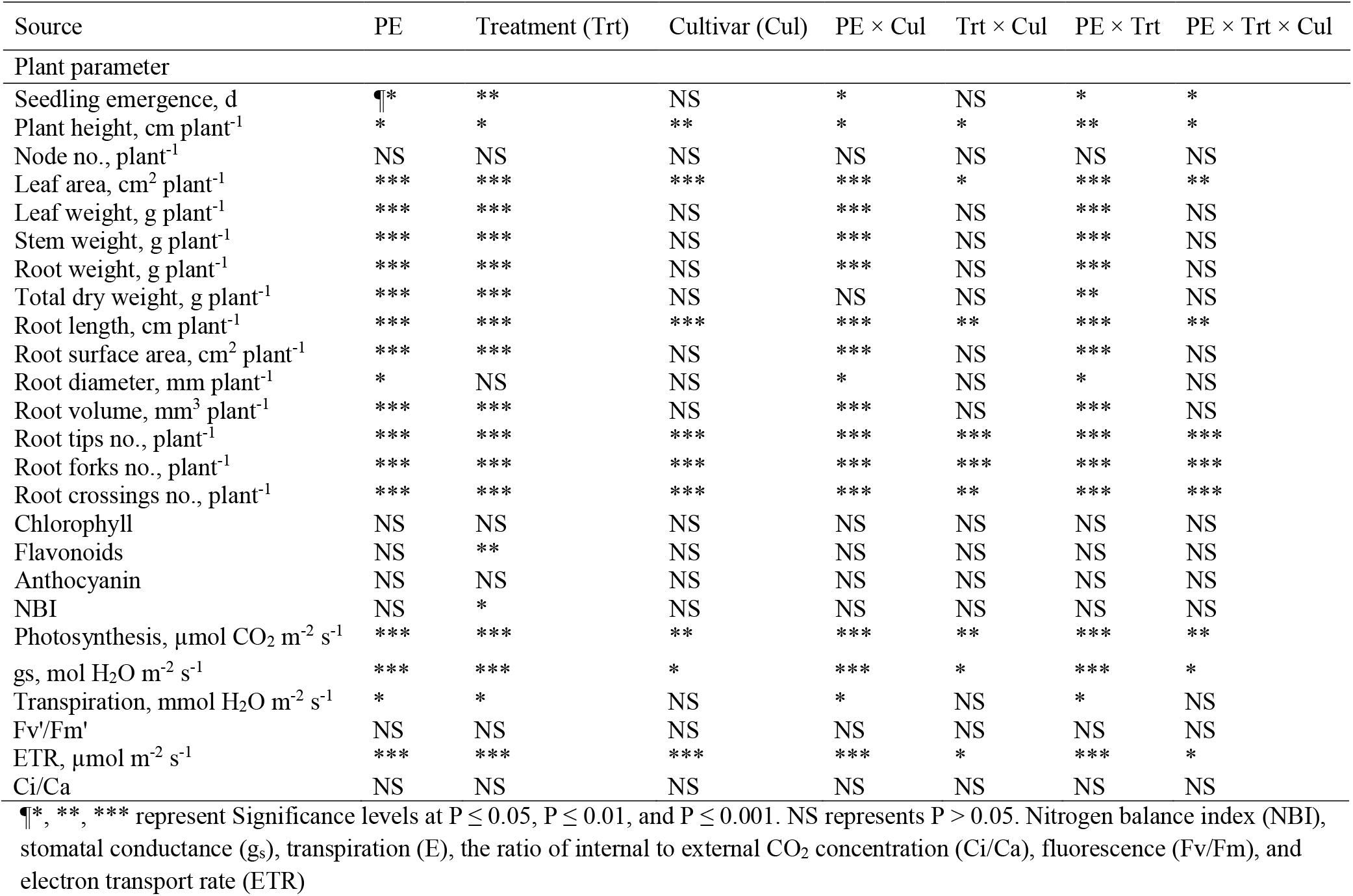
Analysis of variance across the irrigation treatments (Trt), parental environment (PE), cultivars (Cul), and their interaction (Cul × Trt × PE) with soybean vegetative growth, development, physiological, and root traits measured at 18 days after sowing (DAS).

**Fig 9.**
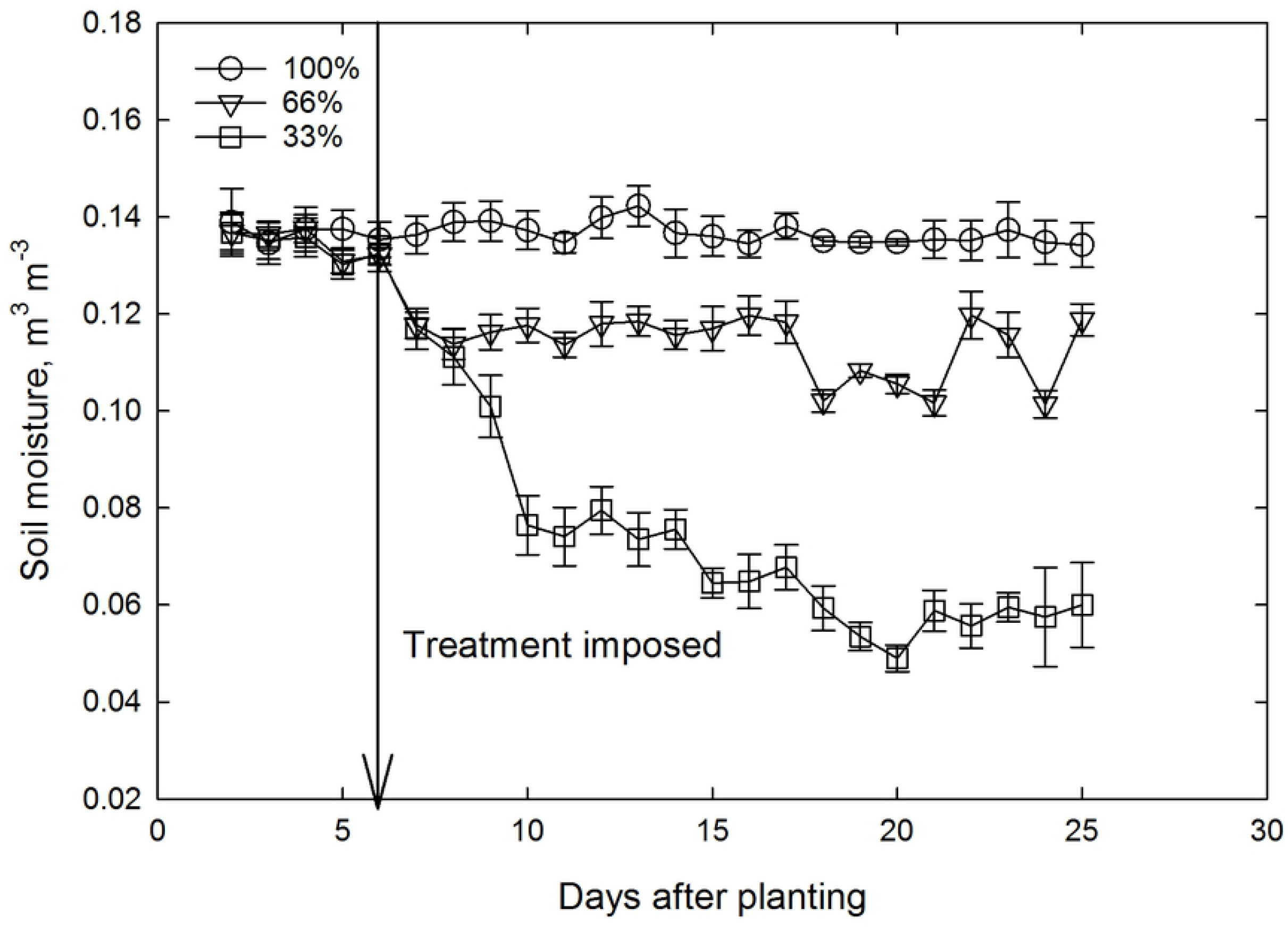
Temporal trends in the average soil moisture content for an experiment where the F1 generation of soybean was exposed to three levels of drought stress starting six days after seeding. Drought stress levels included maintaining the soil moisture content at 100, 66, and 33% of field capacity. The F1 generation was collected from the soybean cultivars Asgrow AG5332 and Progeny P5333RY after exposure to drought stress including replacement of 100, 50, 60, 40, and 20% of the evapotranspiration demand. Symbols represent the mean of eight replications that have been pooled over cultivar. Standard error bars are not displayed when smaller than the symbol for the mean.

**Fig 10.**
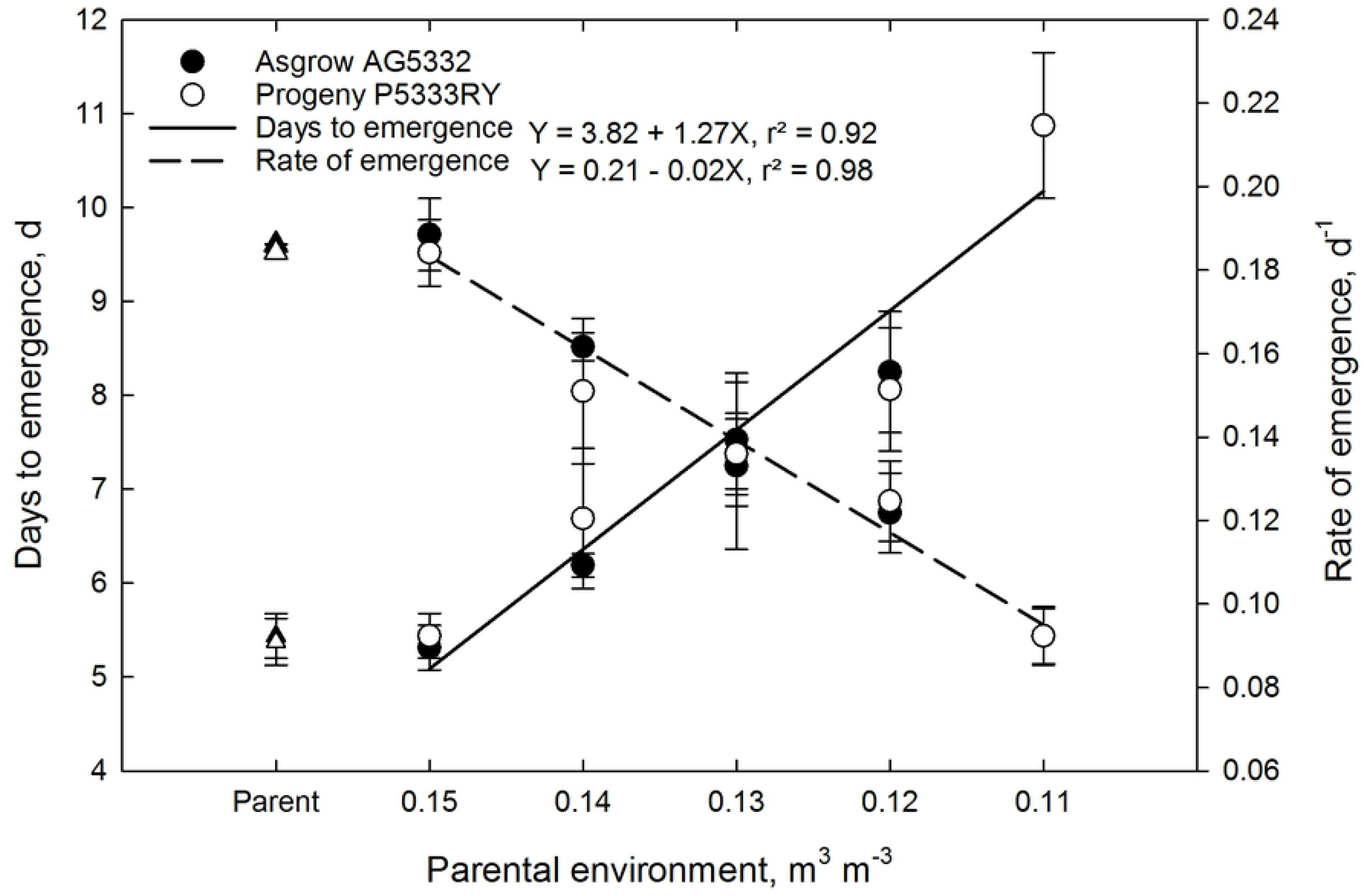
Days to emergence and rate of emergence for the F1 generation of soybean cultivar Asgrow AG5332 and Progeny P5333RY exposed to drought stress including replacement of 100, 80, 60, 40, and 20% of the evapotranspiration demand. Each data point is the mean of four replications pooled over cultivar, while bars represent the standard error of the mean. Lines are the best fit of a linear function.

Drought stress imposed during the reproductive stages of soybean also had a transferable effect on shoot growth and developmental traits (Table 5). Pooled over cultivar, parameters measured at 29 days after seeding, decreased in response to increasing soil moisture stress for plant height (Fig 11), leaf area, and biomass components. The leaf area (Fig 12b), stem and leaf dry weights, and total weight (Fig 12c) of the offspring plants grown under both control (100% FC) and 33% FC were consistently lower than the offspring from control maternal treatment (100% ET) and parents. The mean leaf area and total dry weight were reduced by 48% at the 100% FC (control treatment) in Asgrow AG5332 offspring from 20% ET maternal treatment, compared to the parent plant at the same irrigation treatment. The offspring of Progeny P5333RY from 20% ET maternal treatment, on the other hand, exhibited 61 and 55% reduction for leaf area and total dry weight, respectively, compared to its mother plant (Fig 12).

**Fig 11.**
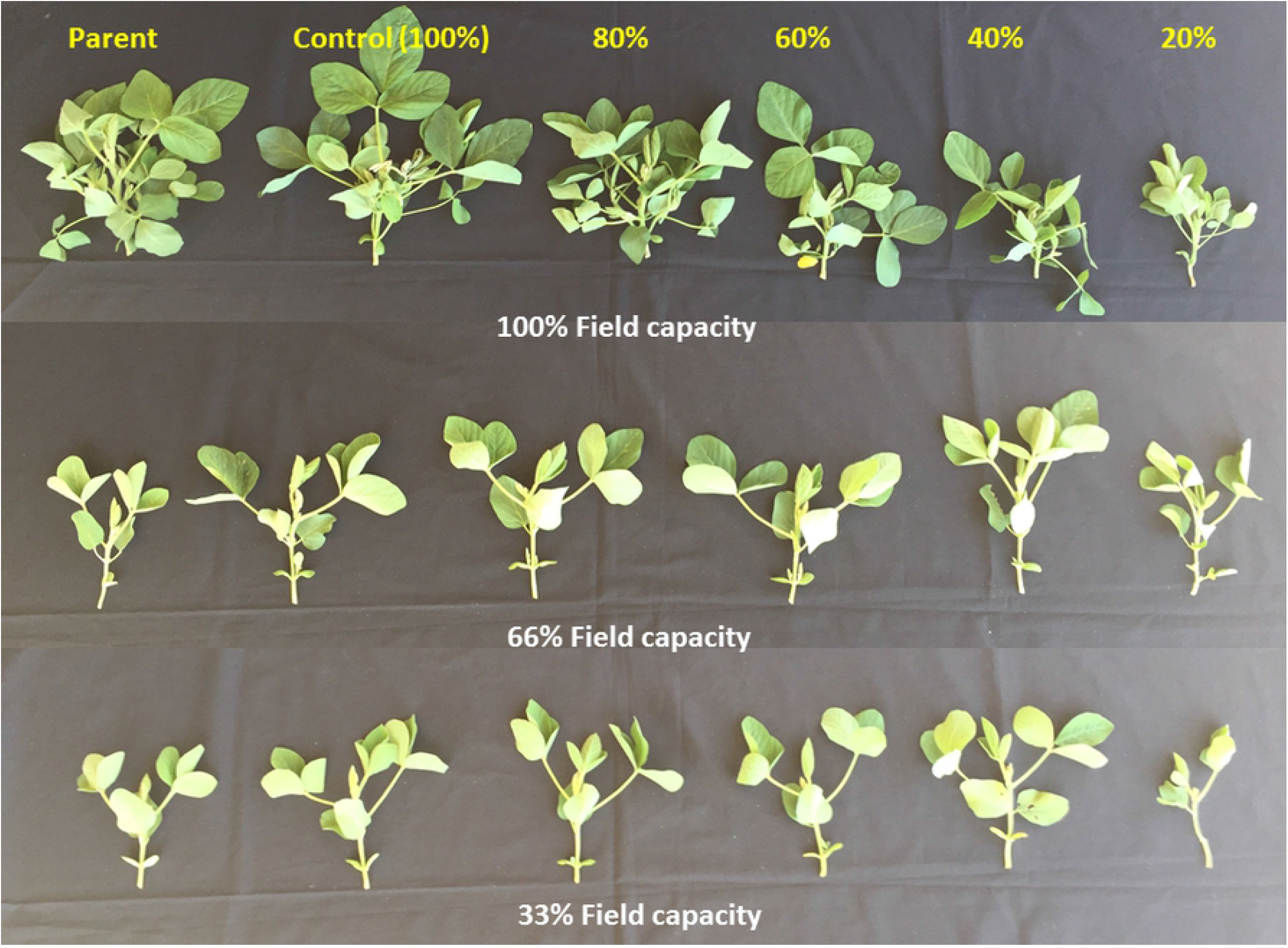
Shoot growth for the F1 generation of soybean cultivar Asgrow AG5332 and Progeny P5333RY exposed to drought stress including replacement of 100, 80, 60, 40, and 20% of the evapotranspiration demand.

**Fig 12.**
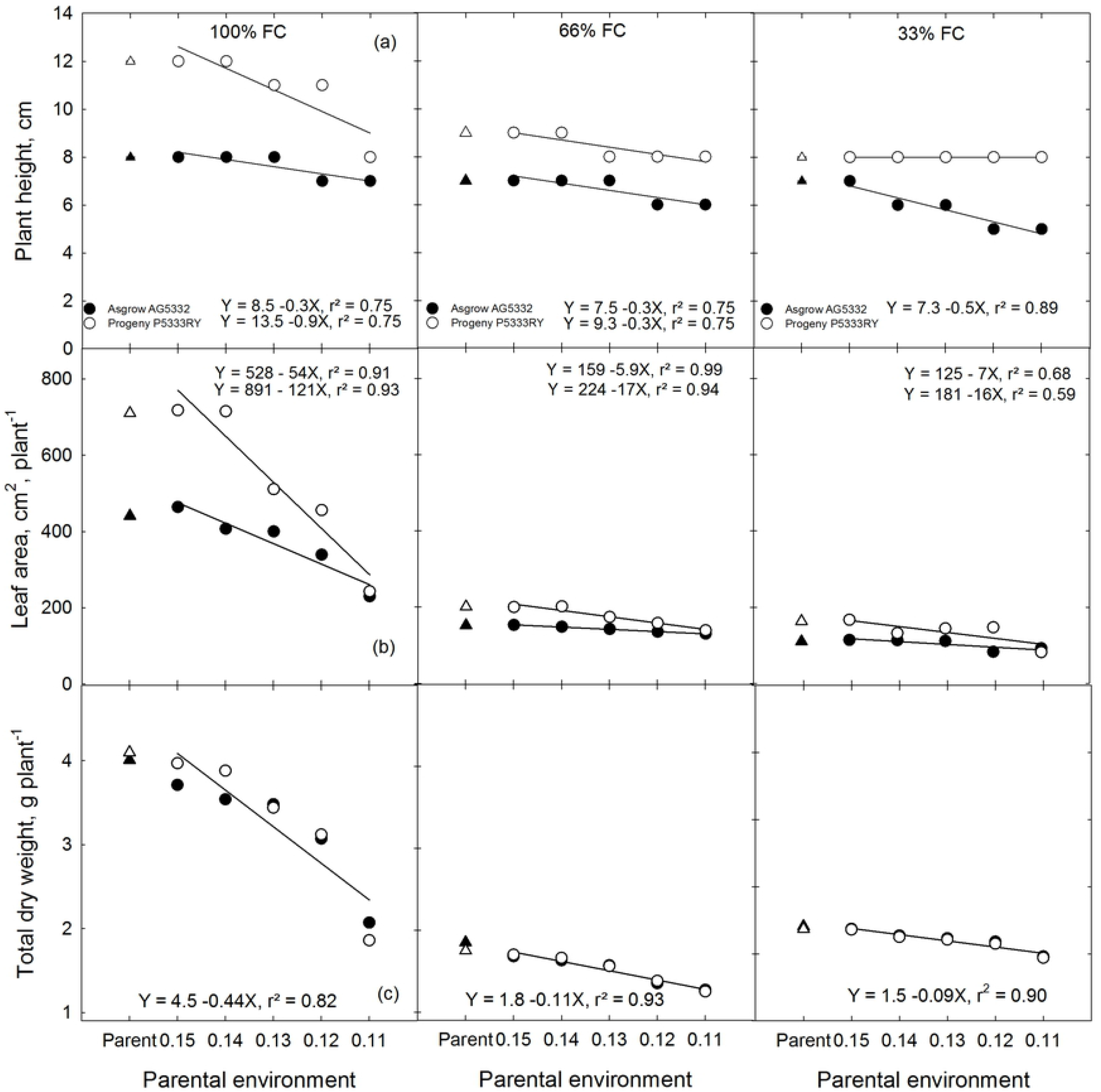
Plant height, leaf area, and total dry weight for the F1 generation of soybean cultivar Asgrow AG5332 and Progeny P5333RY exposed to drought stress including replacement of 100, 80, 60, 40, and 20% of the evapotranspiration demand.

Also, a number of root tips, forks, and crossings were also lower in soil moisture stressed offspring compared to their parents (Fig 13). Regardless the maternal environmental effects, all the soybean plants showed less lateral branching close to the surface layer and more taproot elongation towards the deeper layers of soil under sub-optimal soil moisture conditions (66 and 33% FC), compared to the optimum irrigation (100% FC) (Fig 14). Having a longer tap root system could be a drought-adaptive mechanism to increase water and nutrient uptake under stressed conditions [45,46]. Overall, parental plant and the plant from optimum maternal environment had highly branched, longer, and thicker root system compared to the root systems of offspring plants from stressed maternal environments (Fig 14).

**Fig 13.**
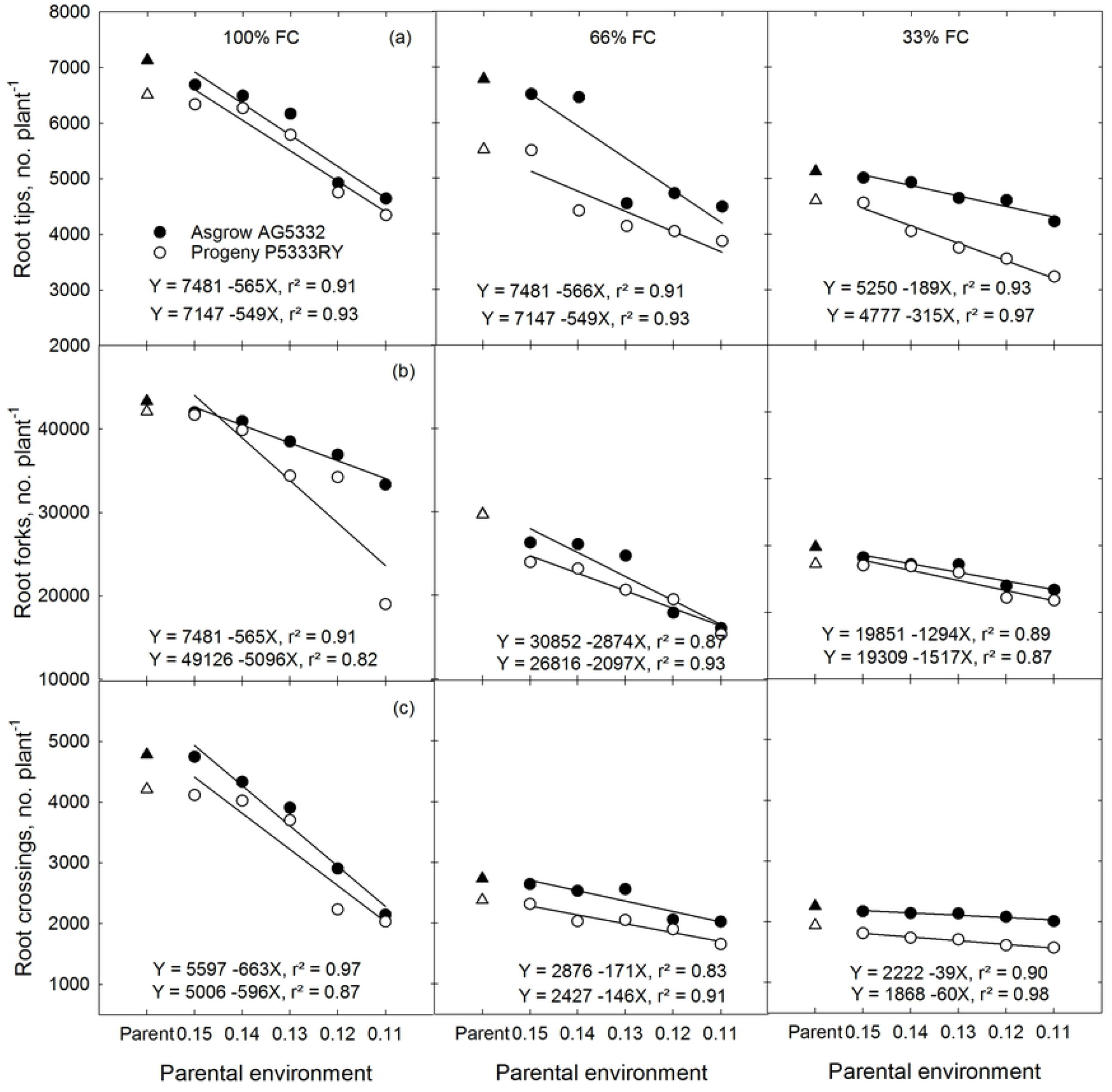
Root tips, forks, and crossings for the F1 generation of soybean cultivar Asgrow AG5332 and Progeny P5333RY exposed to drought stress including replacement of 100, 80, 60, 40, and 20% of the evapotranspiration demand. Each data point is the mean of four replications pooled over cultivar, while bars represent the standard error of the mean. Lines are the best fit of a linear function.

**Fig 14.**
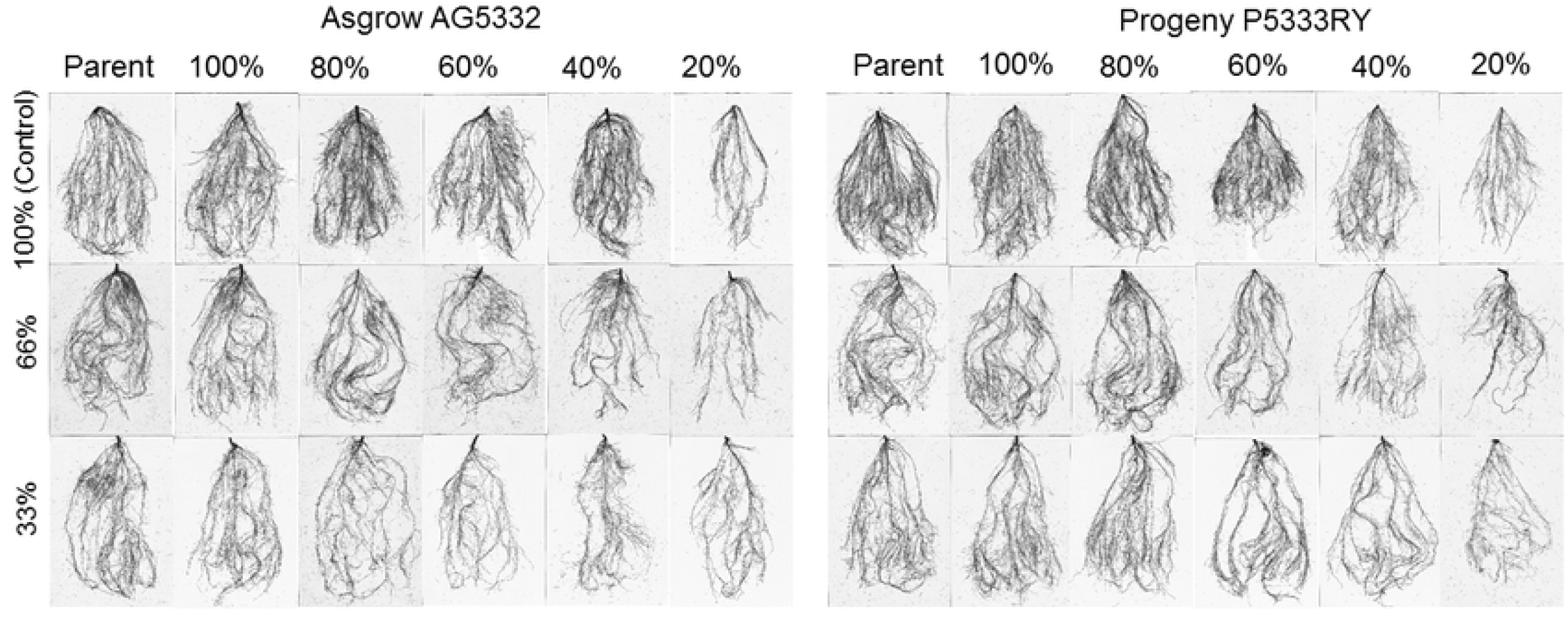
Root growth for the F1 generation of soybean cultivar Asgrow AG5332 and Progeny P5333RY exposed to drought stress including replacement of 100, 80, 60, 40, and 20% of the evapotranspiration demand.

Similar to growth and developmental traits, physiological and gas-exchange traits of the offspring were also affected by maternal environments (Table 5). Lower content of chlorophyll and increased content of flavonoids were observed in offspring from stressed maternal environments (data not presented). Typically, flavonoids production is influenced by genotype and environmental factors [40], particularly an increased production in the leaf as a protective mechanism against abiotic stresses. The net photosynthesis (Pn), stomatal conductance (g_s_), and electron transport rate (ETR) were consistently lower in the offspring plants than the parents irrespective of the irrigation treatment (Fig 15). The Pn was reduced by 62, 57, and 53% in Asgrow AG5332 offspring at 100, 66, and 33% FC while the reduction was 48, 57, and 50% for Progeny P5333RY when compared to their parents (Fig 15a). Regardless of the maternal environments, g_s_ and C_i_/C_a_ both decreased with increasing soil moisture stress. Although many studies have reported that gs is the main factor which is responsible for the net photosynthesis reduction [47,48] compared to non-stomatal limitation such as specific impairments of key metabolic enzymes (Rubisco), decrease in energy consumption, and decrease in the chemical and enzymatic reactions [49,50], our findings suggested the involvement of both stomatal and non-stomatal factors for the decrease in net photosynthesis in soybean both parents and offspring.

**Fig 15.**
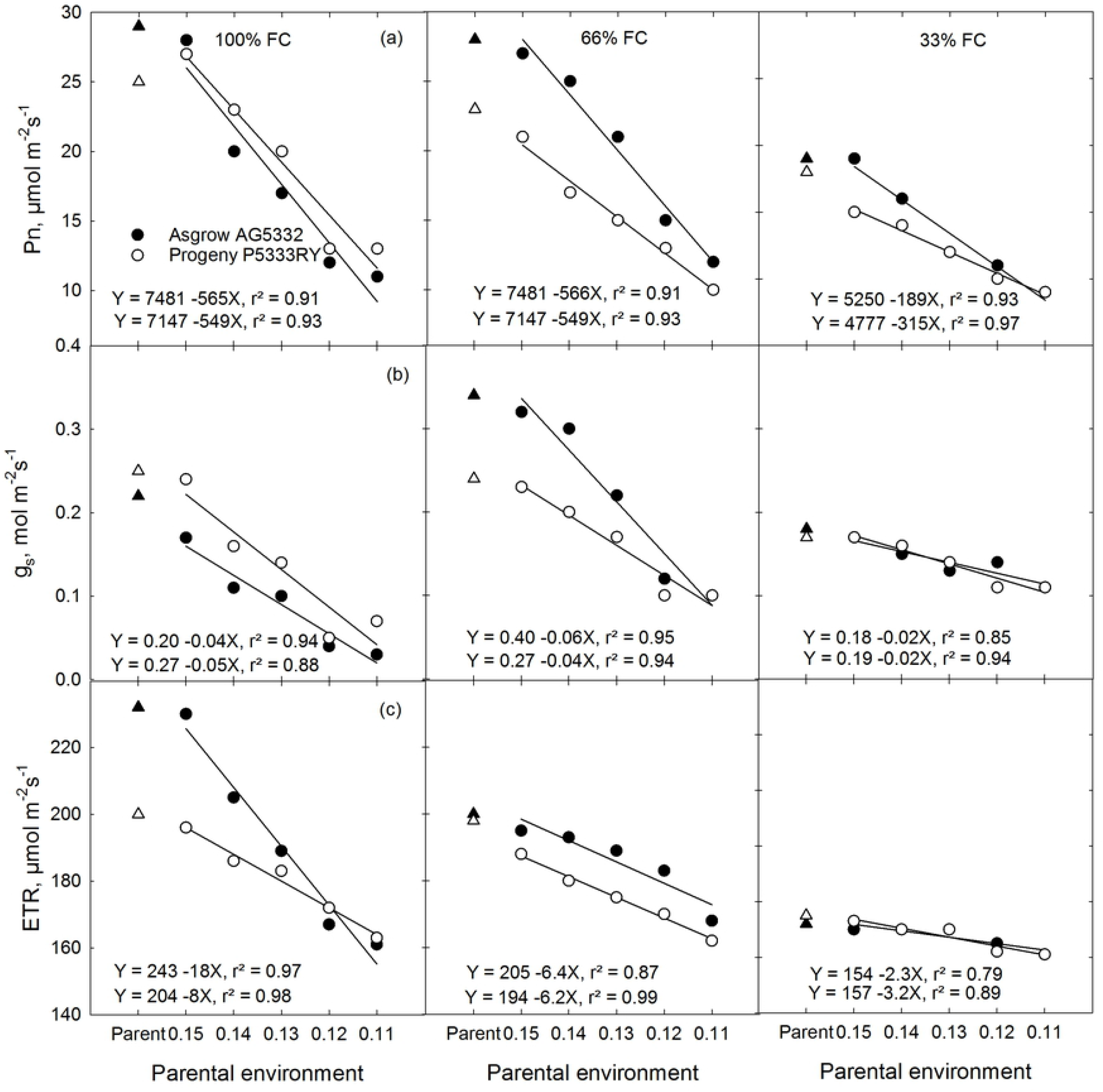
Photosynthesis, stomatal conductance, and electron transport rate for the F1 generation of soybean cultivar Asgrow AG5332 and Progeny P5333RY exposed to drought stress including replacement of 100, 80, 60, 40, and 20% of the evapotranspiration demand. Each data point is the mean of four replications pooled over cultivar, while bars represent the standard error of the mean. Lines are the best fit of a linear function.

In conclusion, this research confirms that drought stress during the reproductive growth of soybean affects the fitness and tolerance of the F1 generation to drought stress. The transgenerational effects of drought stress on the germination of the F1 generation were correlated with seed weight, protein, fatty acids, seed carbohydrates, and minerals. The correlation between germination and the parameters above indicate that the drought stress has an effect on how the maternal lines allocate seed storage reserves, which, subsequently, affects seedling emergence and growth in the F1 generation.

The effect of the environment has serious ramifications for seed production and multiplication companies. Our data indicate that seed produced in environments where drought occurs during or through the reproductive stages is deleterious to seed production and multiplication. The smaller, poor quality seeds that resulted from the stressed maternal environment could result in lower seedling survival in progeny, making them less adapted to withstand drought stress in subsequent generations. However, although we concluded that the main factor responsible for offspring susceptibility to soil moisture stress was the difference in seed size and storage reserve, different epigenesis-based mechanisms also cannot be excluded when describing the maternal effects on offspring fitness.

## Acknowledgments

We thank David Brand for technical assistance and graduate students of the Environmental Plant Physiology Lab at Mississippi State University for their support during data collection. We also thank Sandra Mosley of USDA-ARS, Stoneville, MS, for seed composition analyses. This project was funded by the Mississippi Soybean Promotion Board and the National Institute of Food and Agriculture, 2016-34263-25763 and NIFA-MIS 043040. This article is a contribution from the Department of Plant and Soil Sciences, Mississippi State University, Mississippi Agricultural, and Forestry Experiment Station. Mention of trade names or commercial products in this publication is solely to provide specific information and does not imply recommendation or endorsement by the United States Department of Agriculture (USDA). USDA is an equal opportunity provider and employer.

## Author Contributions

**Conceptualization:** K.R. Reddy, C. Wijewardana

**Data curation:** C. Wijewardana, K.R. Reddy, N. Bellaloui

**Formal analysis:** C. Wijewardana, K.R. Reddy, L.J. Krutz

**Funding acquisition:** K.R. Reddy

**Supervision:** K.R. Reddy

**Methodology:** C. Wijewardana, K.R. Reddy

**Resources:** K.R. Reddy, N. Bellaloui

**Writing-original draft:** C. Wijewardana

**Writing-review & editing:** C. Wijewardana, K.R. Reddy, L.J. Krutz, N. Bellaloui, W. Gao

